# Cilia stimulatory and antibacterial activities of bitter receptor agonist diphenhydramine: insights into potential complimentary strategies for CF nasal infections

**DOI:** 10.1101/2022.01.31.478409

**Authors:** L. E. Kuek, D.B. McMahon, R.Z. Ma, Z.A. Miller, J.F. Jolivert, N.D. Adappa, J.N. Palmer, R.J Lee

## Abstract

**BACKGROUND:** Bitter compounds increase ciliary beating and nitric oxide (NO) production in nasal epithelial cells through T2Rs in motile cilia. We examined expression of cilia T2Rs and both host and bacterial responses to T2R14 agonist diphenhydramine.

**METHOD:** Using cultured human nasal epithelial cells grown at air liquid interface, we measured expression of T2Rs via qPCR. We measured effects of diphenhydramine on ciliary beat frequency via high-speed imaging and nitric oxide production via fluorescent dye DAF-FM. We measured effects of diphenhydramine on growth of lab and clinical strains of *Pseudomonas aeruginosa.* We measured biofilm formation of *P. aeruginosa* using crystal violet staining and surface attachment of *P. aeruginosa* to cystic fibrosis bronchial epithelial (CBFE41o-) cells using CFU counting.

**RESULTS:** T2R expression increased with mucocilliary differentiation and did not vary between CF and non-CF ALIs. Treatment with *P. aeruginosa* flagellin decreased expression of diphenhydramine-responsive T2R14 and 40, among other isoforms. Diphenhydramine increased both NO and CBF. Increases in CBF were disrupted after flagellin treatment. Diphenhydramine impaired growth, biofilm production, and surface attachment of *P. aeruginosa.*

**CONCLUSIONS:** T2R expression is similar between normal and CF cells but decreases with flagellin treatment. Utilizing T2R agonists as therapeutics within the context of CF, *P. aeruginosa* infections may require co-treatment with anti-inflammatories to prevent the reduction of T2R expression with TLR activation. T2R agonist diphenhydramine increases NO production and CBF while also decreasing bacterial growth and biofilm production, and thus diphenhydramine or derivate compounds may have potential clinical usefulness in CF infections as a topical therapy.

**HIGHLIGHTS:**

- T2R14 agonist diphenhydramine increases nitric oxide production and cilia beating
- Flagellin decreases T2R14 expression in primary airway epithelial cells
- T2R14 agonist Diphenhydramine inhibits *Pseudomonas* growth and biofilm formation

## 1 INTRODUCTION

Every inhalation is an opportunity for pathogens to enter the airways. Thus, airway epithelia have multiple protective countermeasures [1, 2]. Mucus secreted by goblet cells entraps pathogens while multi-ciliated cells move pathogens out of the airway via mucociliary clearance (MCC) [3]. MCC is compromised in cystic fibrosis (CF) [4] and other “muco-obstructive” lung diseases [5]. CF is characterized dehydrated ASL and thickened mucus, preventing pathogen removal and leading to respiratory infections (6). Enhancing MCC may boost host defenses in CF. Topical therapies to boost MCC and/or kill bacteria are of particular interest in the nose, where tissue is accessible to high volume nasal rinse (e.g., neti pot) strategies.

Cilia are the first contact points for airway pathogens and are equipped with receptors to recognize these pathogens [3]. Several members of the G protein-coupled bitter taste receptors (taste family 2 receptors or T2Rs) localize to airway cilia [6–9] and can detect bacterial products, including acyl-homoserine lactones [6, 10] and quinolones [11] secreted by CF pathogen *Pseudomonas aeruginosa.* Cilia T2Rs increase ciliary beat frequency (CBF) via calcium-activated nitric oxide (NO) production and PKG phosphorylation of ciliary proteins [12]. The NO produced also has antibacterial effects against *P. aeruginosa* [13]. T2R activation also enhances macrophage phagocytosis [14].

Activating T2Rs may be a strategy to enhance MCC or bactericidal NO-generation in CF-related chronic rhinosinusitis (CF CRS) [15, 16]. CF CRS impairs patient quality of life, but more importantly, CF nasal infections can seed potentially fatal lower respiratory infections [17]. Polymorphisms rendering the cilia-localized T2R38 isoform non-functional are correlated with increased nasal bacterial load, susceptibility to CRS, and worse sinus surgery outcomes in non-CF patients (reviewed in [6]). These same polymorphisms are correlated with more severe nasal symptoms in CF CRS patients [18] and may have relevance to lower airway infections in CF [19].

We sought to examine if diphenhydramine (DPD), a T2R14 agonist [20, 21] and H1 antihistamine, targets T2Rs in nasal cilia to induce NO production and CBF increases. T2R14 is one of the more highly expressed T2R in nasal and lung epithelial cells [8]. Importantly, DPD is water soluble up to the mM range, and thus DPD or derived compounds (e.g., nonabsorbable lectin or dextran conjugates) could be delivered in localized high doses to nasal tissue, which may be useful in a transient stimulation situation as would occur with a high-volume rinse.

We report that T2R expression increases with mucociliary differentiation, but T2R expression is downregulated by toll like receptor 5 (TLR5) agonist flagellin. DPD increases NO production and ciliary beating, but this response is greatly reduced after TLR5 priming. However, DPD reduces *P. aeruginosa* growth and biofilm formation, suggesting potent T2R-independent benefits of DPD.

## 2 MATERIALS AND METHODS

### 2.1 Primary nasal epithelial cell culture

Primary nasal epithelial cell culture was as described [8, 13, 15] in accordance the U.S. Department of Health and Human Services Title 45 CFR 46.116, the Declaration of Helsinki, and University of Pennsylvania guidelines for residual clinical material (Institutional review board #800614). Written informed consent was obtained from patients ≥18 years of age undergoing surgery for CRS or trans-nasal approaches to the skull base for tumor removal. Nasal epithelial cells were obtained through enzymatic dissociation tissue in minimal essential media (MEM; ThermoFisher, Waltham MA) containing 1.4 mg/ml pronase (MilliporeSigma, St. Louis, MO) and 0.1 mg/ml DNase (MilliporeSigma) for 1 hour at 37°C, followed by 2 washes with MEM plus 10% fetal bovine serum. Cells were then incubated on tissue culture plastic in PneumaCult-Ex Plus (Cat.# 05040, Stemcell Technologies, Vancouver BC Canada) containing 100 U/ml penicillin and 100 μg/ml streptomycin (Gibco, Waltham MA) at 37°C, 5% CO2 for 2 hours to allow for adherence and removal of non-epithelial cells (e.g. fibroblasts, macrophages and lymphocytes). Nasal epithelial cells were transferred to a 10 cm tissue culture dish containing PneumaCult-Ex Plus overnight. The following day, culture medium was aspirated and replaced with fresh PneumaCult-Ex Plus. Nasal primary cells were passaged every 4-5 days or when 80% confluent. Cells were seeded for air liquid interface culture (0.33 cm^2^ transwells, 0.4 μm pore size, transparent) as described [8]). Upon apical air exposure, cells were fed every other day with Pneumacult ALI media.

### 2.2 CFBE cell culture

Parental CFBE41o- (not overexpressing Wt or ΔF508 CFTR) were previously provided by D. Greunert (UCSF) and grown in minimal essential media (MEM) plus Earle’s salts containing (ThermoFisher) 1x cell culture pen/strep (Gibco), 1 mM L-glutamine (Gibco), 10% FetalPlex serum substitute (Gemini Biosciences, West Sacramento, CA). Cells were cultured for 5 days on transwells (1 cm^2^ pore size, transparent, Greiner BioOne, Kremsmünster, Austria) before apical aspiration of media and exposure to air. Cells were fed basolaterally for an additional 7 days before use to establish a high transepithelial electrical resistance (TEER) and tight epithelial barrier. ALIs were fed on the basolateral side with the same media used for cell propagation.

### 2.3 Live cell imaging of calcium (Ca^2+^), nitric oxide (NO), production and ciliary beat frequency (CBF)

Imaging of NO and CBF was as described [15]. For NO, ALI cultures were loaded with DAF-FM by incubation in 10 μM DAF-FM diacetate (ThermoFisher) on the apical side in HBSS plus 5 μM carboxy-PTIO (to scavenge baseline NO; Cayman Chemical) for 90 minutes. For submerged CFBEs on 8-well chambered coverglass (CellVis, Sunnyvale, CA), loading was the same but for 45 min instead of 90 min. After washing, imaging was performed using an IX-83 microscope (10x 0.4 NA PlanApo objective, Olympus Life Sciences, Tokyo, Japan) with LED illumination (Excelitas Technologies LED120Boost), 16-bit Orca Flash 4.0 sCMOS camera (Hamamatsu, Japan), standard FITC filter set (470/40 nm excitation, 495 lp dichroic, and 525/40 nm emission; 49002-ET, Chroma Technologies) and MetaFluor (Molecular Devices, Sunnyvale, CA). For cilia beating, cultures were maintained at ~28°C in DPBS (+1.8 mM calcium) on the apical side and HEPES-buffered HBSS supplemented with 1× MEM amino acids (Gibco) on the basolateral side. Sisson-Ammons Video Analysis software was used to measure whole-field CBF. Diphenhydramine and all other reagents used were from Sigma Aldrich unless otherwise specified.

### 2.4 Quantitative PCR

ALIs were lysed in TRIzol (ThermoFisher). RNA was isolated using Direct-zol (Zymo Research). RNA was transcribed to cDNA via High-Capacity cDNA Reverse Transcription Kit (ThermoFisher); cDNA was then quantified utilizing Taqman Q-PCR probes (QuantStudio 5 Real-Time PCR System, ThermoFisher). Data was then analyzed using Microsoft Excel and plotted in GraphPad PRISM.

### 2.5 Bacterial Assays

All bacterial assays growth and biofilm were carried out as described [22]. *Pseudomonas aeruginosa* lab strain PAO-1 was from American Type Culture Collection (HER-1018; ATCC BAA-47). Clinical isolates from *P. aeruginosa* from CF-related CRS were obtained from Dr. Noam Cohen and Dr. Laura Chandler from the Philadelphia VA Medical Center

Department of Otorhinolaryngolgoy and clinical microbiology lab, respectively. Planktonic *P. aeruginosa* or methicillin-resistant *S. aureus* M2 [22] strains were cultured in LB (ThermoFisher) at 37 °C with shaking. For planktonic growth, an overnight log-phase culture was diluted to OD 0.1 in 10 mL total volume per sample. Cultures were grown at 37°C with shaking (180 RPM); 1 mL of solution was removed at each time point and assayed for OD at 600 nm in a spectrophotometer.

*P. aeruginosa* biofilms were grown at 37 °C (no shaking) in 25% LB (low nutrient media to promote biofilm formation) for 5 days in standard 96 well microtiter plates. OD 0.1 cultures were diluted 1:100 in 25% LB and 100 μL was added to each well of a 96 well plate. Plates were sealed with parafilm to prevent dehydration. After incubation, microtiter plates were washed three times with distilled water, followed by staining with 1% crystal violet in water for approximately 30 min. After a second washing, biofilm mass and crystal violet were solubilized by 30% acetic acid for 30 min with gentle occasional shaking, and read on a plate reader (Spark 10M, Tecan) at 590 nm.

Bacterial attachment assays were carried out similarly to [23]. CBFE ALI cultures (12-well plate size, 1 cm^2^ surface area) were washed on the basolateral side with PBS and transferred into antibiotic-free media 48 hrs before use in these assays. The apical side was washed three times with PBS before use. *P. aeruginosa* were grown overnight, diluted to OD 0.1 in sterile serum-free DMEM with shaking for 1 hour at 37°C. CFBE ALIs were inoculated on the apical side with 3×10^6^ CFU in 0.5 ml in media ± diphenhydramine ± ceterizine. Co-cultures were incubated for 1 hour at 37°C, followed by washing the apical side 3x with sterile media and a further 4 hour incubation with 500 μL sterile media on the apical side. Cultures were then washed again with sterile media on the apical side. Remaining adherent bacteria were removed by washing with sterile PBS + 1% triton X-100. This solution was then serially diluted 3 times and 10 μL was spotted on LB plates for CFU counting.

### 2.6 Immunofluorescence microscopy

ALI cultures were fixed in 4% paraformaldehyde for 20 min at room temperature. Blocking and permeabilization was in DPBS + 5% normal donkey serum, 1% BSA, 0.2% saponin, and 0.3% Triton X-100 for 45 min at 4°C. Cultures were incubated in primary antibody at 1:100 in blocking buffer (no triton) at 4°C overnight. Cultures were then incubated with AlexaFluor-labeled donkey anti-mouse or anti-rabbit (1:1000) at 4°C for 1 hour. ALIs were cut out and mounted with Fluoroshield with DAPI mounting media (Abcam, Cambridge, MA). Images were taken on an Olympus IX-83 microscope using x60 objective (1.4 NA oil; MetaMorph software). T2R14 (PA5-39710) was from ThermoFisher.

Submerged CFBEs were cultured on collagen-coated glass (MatTek, Ashland MA) and fixed in 3.2% paraformaldehyde for 10 min followed by blocking/permeabilization in DPBS + 5% normal donkey serum, 1% BSA, 0.2% saponin, and 0.1% Triton X-100 for 30 min at 4°C. Primary antibodies used were for GLUT-1 (mouse monoclonal SPM498; MS-10637-P0; LabVision, San Francisco, CA), endothelial (e) NOS (NOS3; SC-654 rabbit polyclonal, SantaCruz Biotechnology, Dallas TX), inducible (i) NOS (NOS2; NB300-605S5), and neuronal (n) NOS (NOS1; ab5586)

### 2.7 Data analysis and statistics

Analyses were performed in Excel or GraphPad Prism; *p* <0.05 was considered statistically significant. One-way analysis of variance (ANOVA) was performed with Dunnett’s posttest (comparisons to control group), Tukey-Kramer posttest (for comparisons of all samples), or Bonferroni posttest (preselected pair-wise comparisons). All bar graphs are mean ± SEM; * indicates *p* < 0.05 and ** indicates *p* <0.01. Images were analyzed in ImageJ (W. Rasband, NIH, Bethesda MD).

## 3 RESULTS

### 3.1 T2R expression increases with nasal mucociliary differentiation but is decreased by P. aeruginosa flagellin

Upon air exposure, cultured primary human nasal epithelial basal cells differentiate to ciliated cells which contain several bitter taste receptor isoforms (T2Rs) that act as sentinels in the innate immune system. Differentiation of non-CF nasal air-liquid interface cultures (ALIs) was determined by the increase in cilia marker MS4A8B [3, 24], which increased to a maximal after 3-4 weeks of exposure to air (**Figure 1A-B**). Transcripts for cilia-localized T2R’s, including T2R14, increased over the same time course in non-CF ALIs (**Figure 1A**). No significant difference was observed between T2R transcript levels in non-CF vs F508del/F508del homozygous CF ALIs after differentiation (**Figure 1C**) at week 21. This fits with published gene array data suggesting T2R expression is similar between non-CF and CF cells (**Supplementary Figure 1**).

**Fig. 1.**
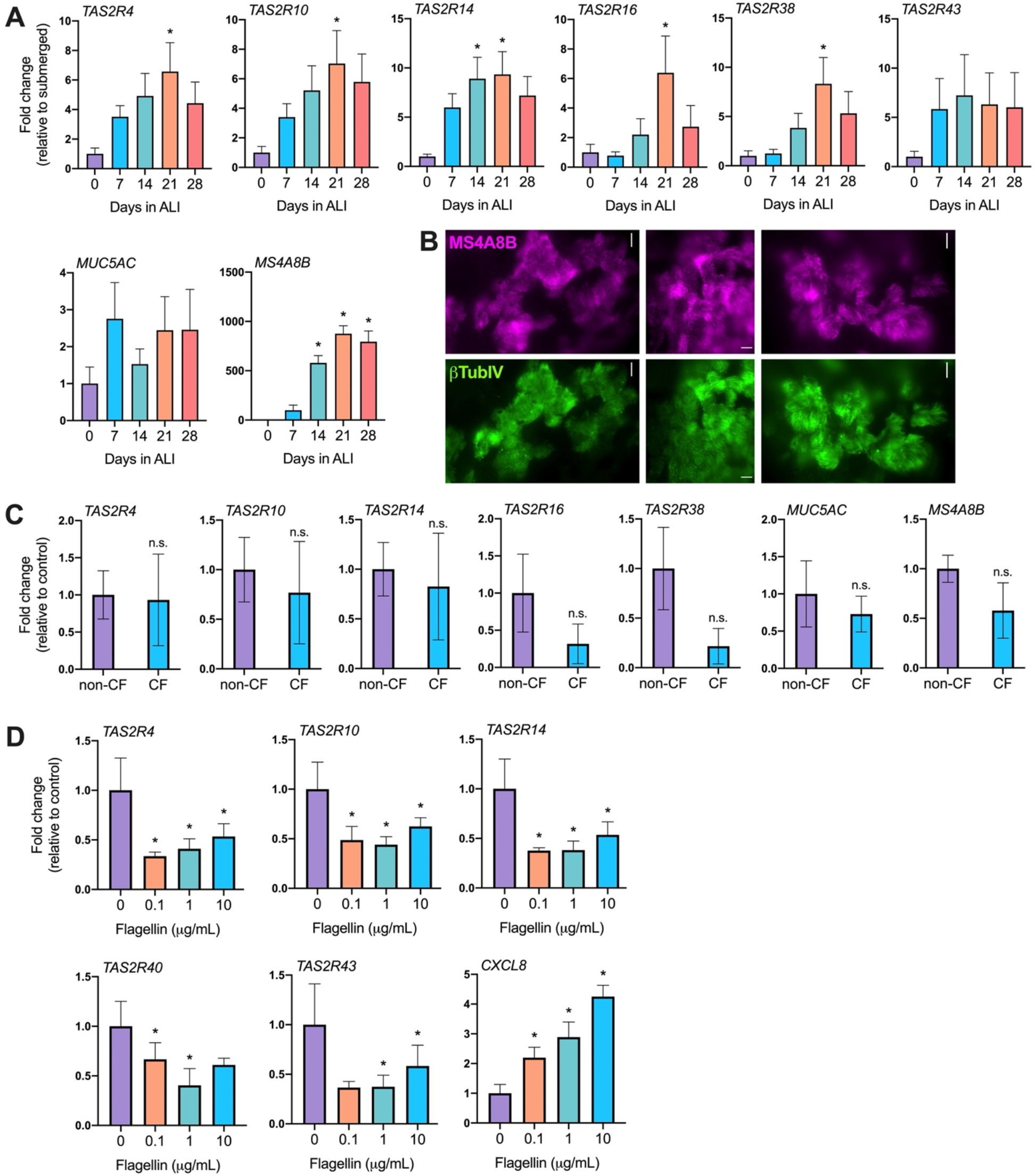
T2R expression in primary nasal air liquid interface (ALI) cultures. **(A)** T2R expression increases during differentiation of nasal ALIs. Ciliary T2Rs, 4, 10, 14, 16, and 38 Increase mRNA expression with at least 3 weeks of differentiation as revealed by qPCR. Data points represent the average of 5 patients; significance was analyzed by one-way ANOVA using Dunnett’s posttest (* *p*<0.05). **(B)** Immunofluorescence of MS4A8B (Mouse monoclonal antibody 3E6, Kerafast, Boston, MA) and ß-tubulin IV (Rabbit monoclonal EPR16775, Abcam) showing cilia co-localization. Representative images shown from three ALIs from 3 individual patients. **(C)** Cystic Fibrosis does not alter T2R mRNA Expression. Cilia-localized T2Rs 4, 10, 14, 16, and 38 did not significantly alter expression when comparing nasal ALIs derived from either non-CF or CF patients. F-G, Cilia marker MS4A8B and MUC5AC expression demonstrates that cultures were differentiated. Bar graphs represent data obtained from 5 non-CF and 3 CF patients. Data pairs were analyzed by Students t-test showing no significant differences. **(D)** Flagellin Decreases T2R Expression in nasal ALIs. Human nasal epithelial cells were isolated and grown in ALI cultures then exposed to air to induce differentiation for at least 4 weeks prior to treatment. A-F, Cultures were treated with various concentrations of flagellin (0.1, 1, or 10 μg/mL) for 24 hours prior to RNA analysis via qPCR. Bar graphs represent combined data from 4 patients. Data was analyzed by one-way ANOVA, Dunnett’s posttest; * *p*<0.05.

This initially suggested that T2Rs may be therapeutic targets in CF nasal cells as proposed in non-CF nasal cells [6]. However, we also surprisingly found that *P. aeruginosa* flagellin, a TLR5 agonist [25], reduced expression of T2R14 and other T2R isoforms (**Figure 1D**). Differentiated nasal ALIs were treated with media containing 0.1, 1, or 10 μg/mL of flagellin overnight. Exposure to flagellin downregulated T2R4, 10, 14, 40, and 43 transcripts while IL-8 (CXCL8) expression increased, as expected (**Figure 1D**).

### 3.2 Diphenhydramine activates NO production and CBF in differentiated airway epithelial cells

As described above, T2Rs in cilia activate NO production [6]. We tested if NO production is activated by diphenhydramine, a commonly used H1 antihistamine that is also a T2R14 and 40 agonist [21]. T2R14 is expressed in both bronchial [7] and nasal [15] cilia (**Figure 2A**). T2R14 is a receptor activated by a broad range of pharmaceutical drugs with established safety data [26] as well as plant products used in homeopathic remedies [15, 21]. The wealth of T2R14 agonists means this receptor might be more easily exploited to activate innate immune responses, particularly if we can find agonists that also have antibacterial effects. We hypothesized that finding an agonist with high solubility that could be delivered at high doses might enhance any antibacterial effects.

**Fig. 2.**
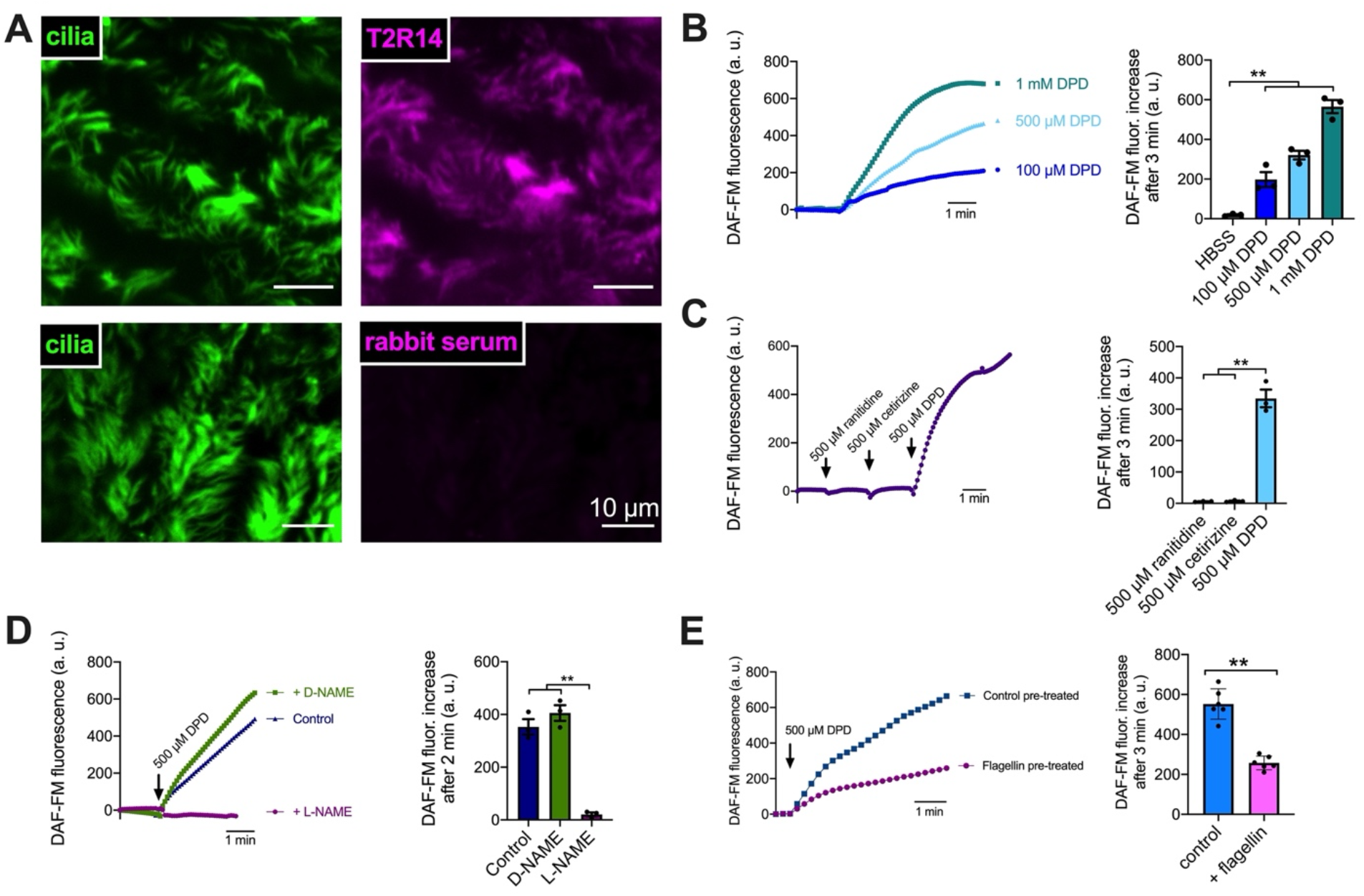
Diphenhydramine (DPD) Induces Nitric Oxide (NO) Production in nasal ALIs. **(A)** T2R14 localizes to cilia in nasal ALIs. Representative immunofluorescence image of nasal cilia (ß-tubulin IV; green) and T2R14 (magenta) from ALI cultures from two individual donors. No staining was observed with rabbit serum plus secondary control (bottom). **(B)** NO production, as revealed by DAF-FM fluorescence, increased in a dose-dependent manner with DPD. **(C)** Antihistamines ranitidine and cetirizine did not initiate NO production, while equimolar diphenhydramine did. **(D)** NOS inhibitor L-NAME (10 μM), blocked NO production while negative control D-NAME (10 μM) did not. **(E)** Flagellin pre-treatment resulted in less NO production in response to DPD. Representative traces from ≥3 experiments using ALIs from different non-CF patients are shown. Bar graphs are mean ± SEM analyzed via one-way ANOVA (B-D) or Student’s *t* test (E); \*\**p*<0.01.

We tested diphenhydramine (DPD), a T2R14 agonist more commonly known as a first generation H1 antihistamine. As expected from a T2R agonist, DPD (100 μM-1 mM) activated NO production in nasal ALIs as visualized via NO-sensitive dye DAF-FM fluorescence, (**Figure 2B**). Equimolar H1 or H2 antagonists cetirizine or ranitidine, respectively, which are not known to activate T2R14, did not activate NO production (**Figure 2C**). Thus, this is likely an effect of T2R stimulation rather than H1 antagonism. NO synthase (NOS) inhibitor L-NAME (10 μM, 1hr pretreatment) inhibited DPD-induced NO production, while negative control D-NAME did not (**Figure 2D**). Fitting with flagellin lowering T2R14 transcript (**Figure 1C**), when primary nasal ALIs were treated with flagellin (0.1 μg/mL) overnight, the DAF-FM fluorescence responses were reduced by ~50% (**Figure 2E**). We also saw Ca^2+^ and NO responses with DPD in CFBE41o- (CFBE) cells, which expressed both T2R14 and eNOS (**Supplementary Figure 2**).

T2R NO production increases CBF [6]. DPD (100 μM) increased CBF by ~1 Hz (~10%; **Figure 3A**), comparable with T2R-induced CBF increases in other studies [13, 15]. Overnight stimulation with flagellin (0.1 μg/mL) caused a loss of the DPD-induced CBF increase (**Figure 3B**). These results suggest DPD CBF increases are due to T2R activation. Moreover, they suggest that while T2Rs can be activated for beneficial NO and CBF responses in CF cells, other inflammatory factors released by *P. aeruginosa* may limit T2R responses by downregulating T2R expression.

**Fig. 3.**
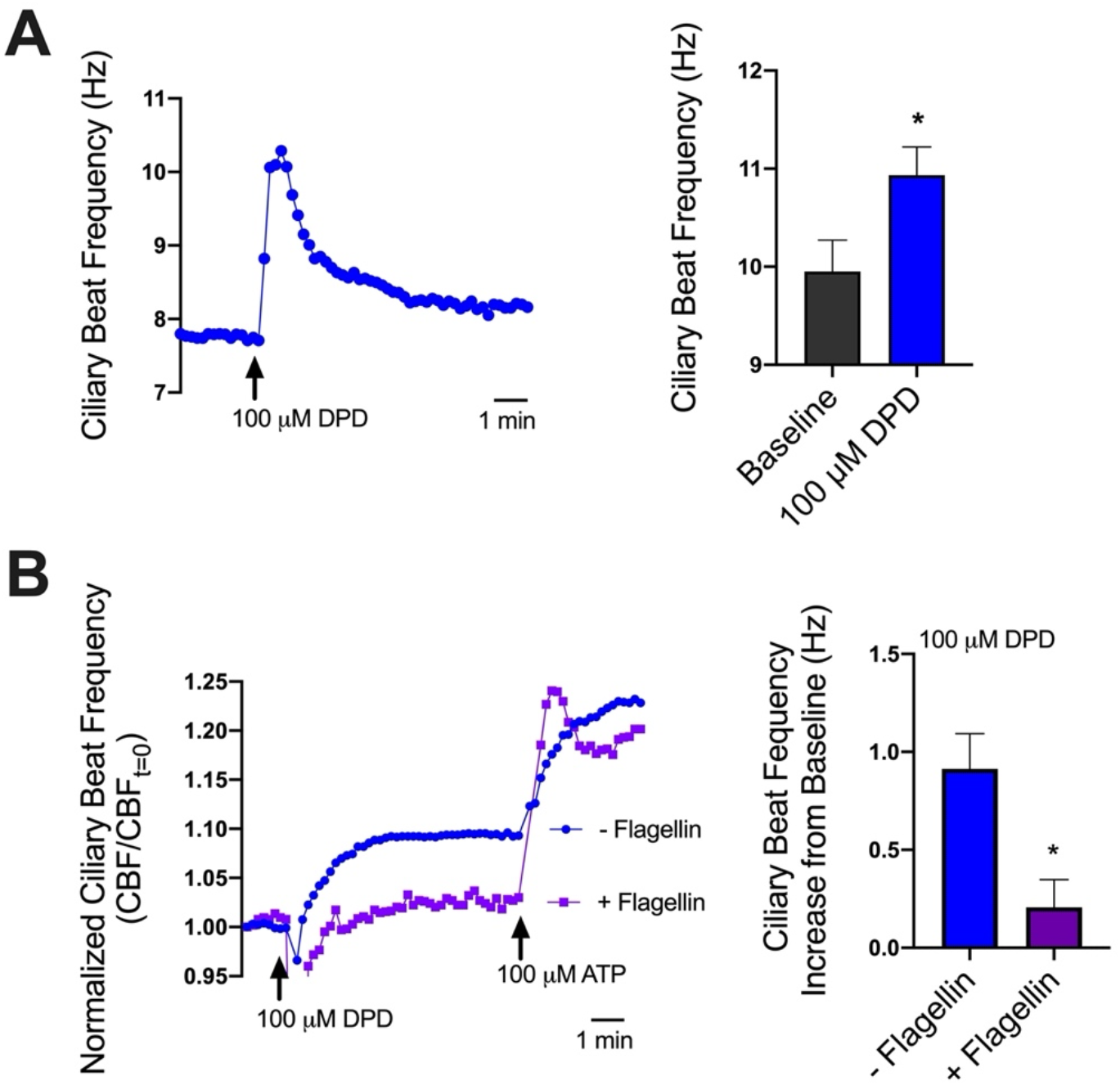
Flagellin Represses DPD Enhancement of Ciliary Beat Frequency. **(A)** T2R14 agonist diphenhydramine (100 μM) increases CBF by up to 1 Hz. **(B)** Fully differentiated (day 21) nasal ALI cultures were treated with media containing 0.1 μg/mL flagellin for 72 hours to ensure reduction in both RNA and protein expression. Flagellin treated cultures did not reveal a 1 Hz increase in CBF when treated with diphenhydramine. Traces are from 1 patient representative of results from ≥3 patients. Bar graphs are combined data from a ≥3 ALIs from ≥3 individual patients.

### 3.3 Diphenhydramine inhibits P. aeruginosa growth, biofilm production, and attachment to CFBE cells

DPD inhibited planktonic growth of both *Staphylococcus aureus* (methicillin-resistant lab strain M2; Figure 4A) and *P. aeruginosa* (PAO1; **Figure 4B**) respectively. We further explored *P. aeruginosa* because of its relevance to CF as well as the larger effect of DPD on *P. aeruginosa* compared with *S. aureus.* DPD also reduced planktonic growth of clinical strains of *P. aeruginosa* (**Figure 4C-G**).

**Fig. 4.**
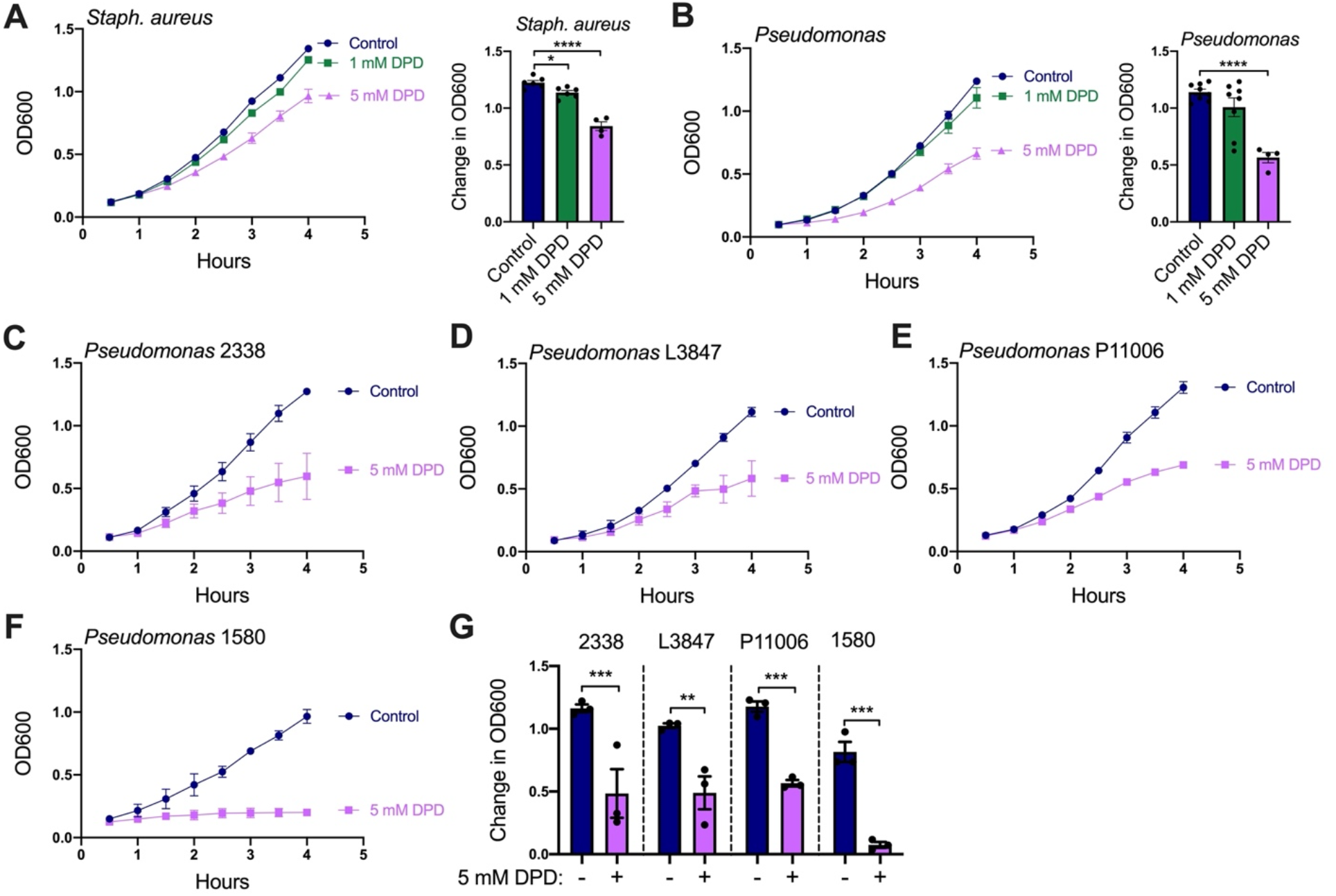
Diphenhydramine Represses the Planktonic Growth of *S. aureus* and *P. aeruginosa.* **(A)** *S. aureus* strain M2 growth (measured by OD_600_) was reduced by 1 to 5 mM diphenhydramine. **(B)** DPD (5 mM) significantly decreased the growth of *Pseudomonas* lab strain PAO-1. **(C-f)** The growth of clinical strains of *Pseudomonas* 2338 (C), L3847 (D), P11006 (E), and 1580 (F) were all impaired by 5 mM DPD. **(G)** Bar graph summarizing results from C-F. Traces are representative of ≥3 experiments with mean ± SEM in bar graphs analyzed via oneway ANOVA using Sidak’s post-test (\*\**p*<0.01, \*\*\**p*<0.001, \*\*\*\**p*<0.0001).

DPD (1-5 mM) also inhibited biofilm production of *P. aeruginosa* PAO1 and clinical CRS *P. aeruginosa* isolate P11006, as visualized through crystal violet staining (**Figure 5A-B**). We tested if this affected *P. aeruginosa* binding to CFBE cells. CFBE cells were incubated with several lab strains of *P. aeruginosa* and P11006 ± DPD on the apical side, followed by washing and recovery of adherent bacteria, which were quantified by CFU counting. DPD caused 5-10-fold less bacteria to adhere to the CFBE cells (**Figure 5C-D**). No toxic effects of DPD were observed in CFBE ALIs by visual inspection of the cultures (**Figure 5E**) or changes in transepithelial resistance (**Figure 5F**).

**Fig. 5.**
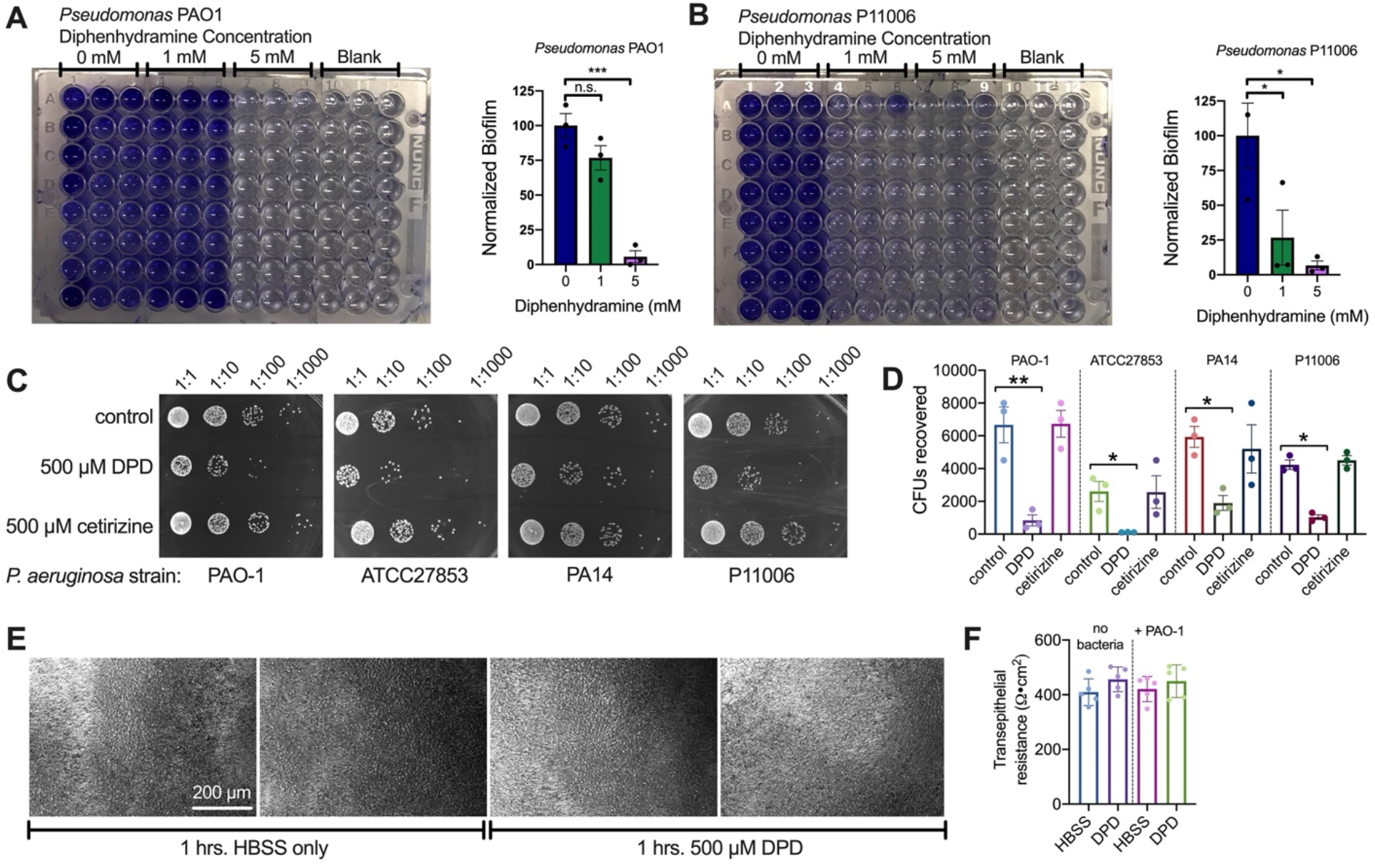
Diphenhydramine (DPD) Inhibits *P. aeruginosa* Biofilm Production and Attachment to CFBE cells. **(A-B)** Representative images of crystal violet staining showing diphenhydramine reduced biofilm mass at 5 mM for lab strain PAO1 (A) and 1 mM for clinical strain P11006 (B). Other clinical strains used in Fig 4 did not form biofilms on 96 well plates and so could not be used in this assay. Representative images of 96-well plate are shown. Data from 3 experiments are combined in bar graphs analyzed by one-way ANOVA using Dunnett’s posttest (* *p*<0.05, *** *p*<0.001). **(C)** Representative images of CFU counts following CFBE attachment assasy as described in the text. DPD (500 μM) reduced the amount of bacteria recovered from CFBE cells while equimolar cetirizine did not. Control is media only. Lab strains PAO-1, ATCC27853, and PA-14 and clinical strain P11006 were used. **(D)** Quantification of results from independent experiments as in C. Significance by one way ANOVA with Bonferroni posttest comparing all values to respective control.; *p<0.5, **p<0.01. **(E)** Representative phase contrast images of CFBE ALIs incubated in HBSS ± 500 μM DPD (1 hour; 37° C). No signs of epithelial break down were observed. Representative of 6 ALIs per condition from independent experiments. F, Transepithelial resistance was measured ±DPD stimulation as in E. No significance difference was observed by one way ANOVA.

## 4 DISCUSSION

Targeting cilia T2Rs may activate CBF and NO-dependent bacterial killing in CF-related CRS or non-CF CRS patients (20). The endogenous agonists known to activate airway cilia T2Rs (acyl-homoserine lactones [13] and quinolones [11]) are produced by *P. aeruginosa.* We found that *P. aeruginosa* are more sensitive to NO-dependent killing than *Staphylococcus aureus* [27].

This suggests that the cilia T2Rs may have evolved to detect the bacteria most susceptible to this defensive pathway. Further work is needed to clarify interactions of specific bacterial species with cilia T2Rs.

We found that T2R expression is similar between CF and non-CF patients in the absence of inflammatory stimuli. However, inflammation in CF may impair signaling of some T2Rs. We saw decreased T2R14 expression and NO/cilia responses to T2R14 agonist diphenhydramine after *Pseudomonas* flagellin pretreatment. This was surprising; we initially expected an increase in T2R expression since T2Rs are tied to innate defense. NFkB signaling downstream of TLR5 may shifts the main defensive pathway toward increased production of antimicrobial peptides like defensins or enhanced NO production via inducible (i)NOS.

DPD, a T2R14 agonist in addition to H1 antihistamine, induces T2R innate immune responses, increasing CBF and NO production. DPD also disrupts *P. aeruginosa* growth, biofilm production, and cell attachment of both lab and clinical strains. DPD was shown to have antibacterial effects against *S. aureus* [28] and may potentiate effects of levofloxacin against *S. aureus* or *P. aeruginosa* [29]. We show effects of DPD alone against *P. aeruginosa* at concentrations that activate T2R receptors. We previously showed antibacterial activity of other T2R14 agonist plant flavones [22]. While DPD may not be the most optimal T2R agonist to use clinically due to the drowsiness induced by first generation H1 antihistamines [30], it has a much higher water solubility than flavones and is already widely used. Alternatively, a DPD-related compound and/or a modified non-absorbable form of DPD (e.g., a dextran or lectin conjugate) that maintains both bitterness and anti-bacterial effects might be a useful to include in a nasal rinse to treat infections. While the mechanism for DPD’s effect on bacteria must be investigated in future work, these observations suggest potential dual action in the airway by activating the host’s innate immunity and decreasing bacterial growth and biofilm formation. Our data also suggest that finding other T2R agonists with antibacterial effects is likely.

## 5 CONCLUSIONS

DPD increased both CBF and NO production in nasal ALIs, consistent with T2R activation. However, the responses to DPD were inhibited by priming of cells with *P. aeruginosa* flagellin, likely due to TLR5 signaling. The full benefits of T2R activation may require co-stimulation with an anti-inflammatories to reduce TLR signaling within the context of CF inflammation to enhance T2R expression. However, DPD inhibits *P. aeruginosa* growth, biofilm production, and attachment to lung epithelial cells through unknown mechanisms. This dual action of DPD may have therapeutic applications in CF patient nasal infections.

## ACKNOWLEDGEMENTS

The authors thank M. Victoria for excellent technical assistance.

## ABBREVIATIONS

ALI: air liquid interface
CBF: ciliary beat frequency
CF: cystic fibrosis
CFBE: cystic fibrosis bronchial epithelial cell
CRS: chronic rhinosinusitis
DPD: diphenhydramine
eNOS: endothelial nitric oxide synthase
GPCR: G protein-coupled receptor
MCC: muciliary clearance
NO: nitric oxide
NOS: nitric oxide synthase
T2R: taste family 2 bitter GPCR
TAS2R: encoding T2R receptor
TLR: toll like receptor

