## Supplementary Figures for "Cilia stimulatory and antibacterial activities of bitter receptor agonist diphenhydramine: insights into potential complimentary strategies for CF nasal infections"

**Supplementary Data:** Kuek, *et al.* Immunomodulatory effects of diphenhydramine.

### Supplementary Figure 1

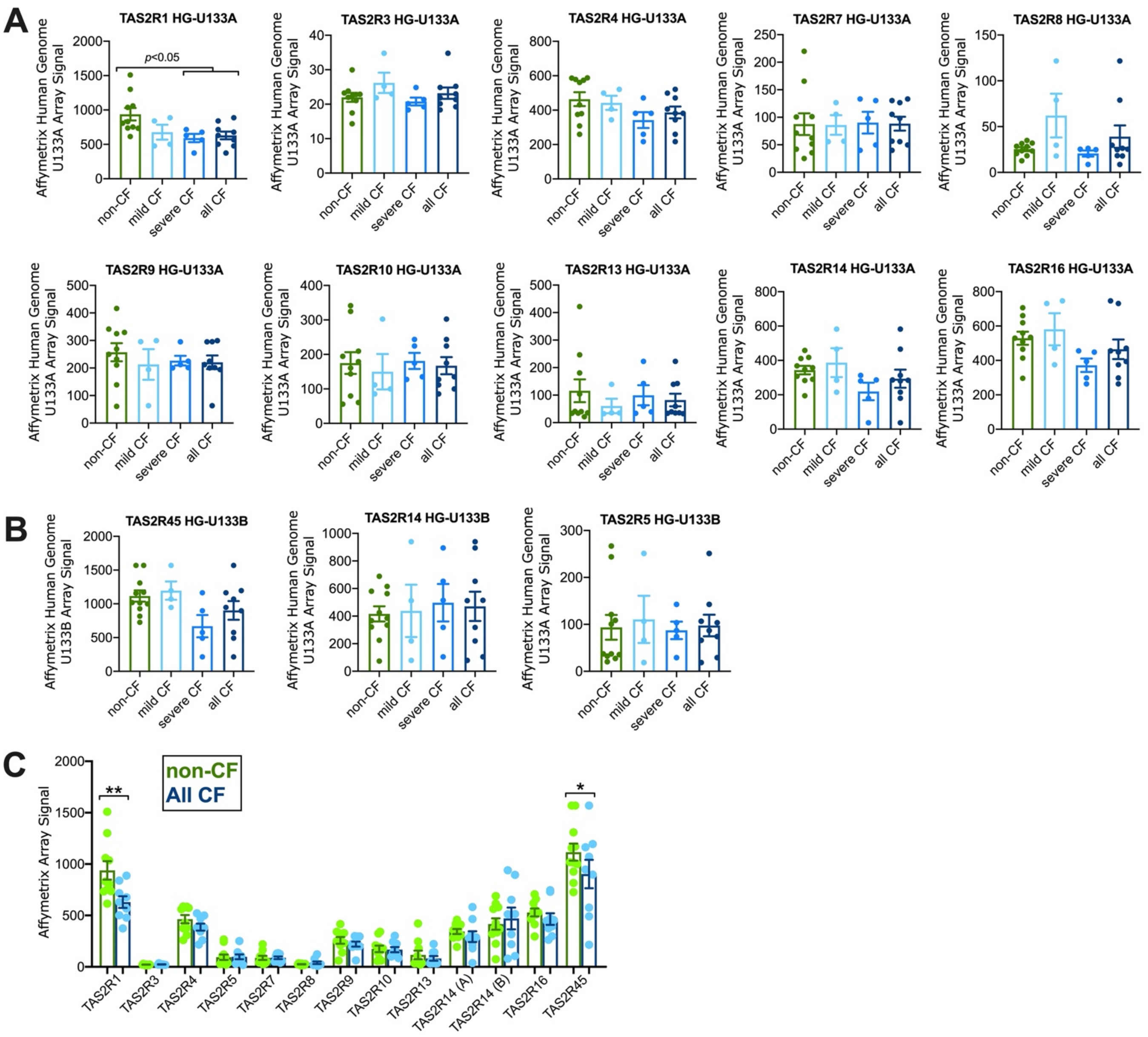

**Supplementary Figure 1: TAS2R gene expression in non-CF vs CF nasal epithelium.** Previously analyzed Affymetrix Human Genome U133A and U133B array expression data from [1] was downloaded from NCBI Gene Expression Omnibus (Reference Series GSE2395, data set records GDS2142 and GDS2143). These data analyzed nasal epithelium of non-cystic fibrosis (CF) vs cystic fibrosis (CF) patients with mild or severe lung disease as defined by the investigators. CF patients were homozygous F508del CFTR. **(A-B)** Changes in expression of TAS2R genes encoding individual T2R receptors that were contained in these arrays (HG-U133A array shown in *A* and HG-U133B array shown in *B*) are plotted and analyzed by one-way ANOVA with Dunnett's posttest comparing all values to non-CF. No significant differences were observed except for a decrease in TAS2R1 encoding T2R1 in severe CF and all CF groups. The all CF group is the pool of mild + severe CF groups. Notably, no change in T2R14 was observed on either HG-U133A or HG-U133B array. **(C)** The relative expression of the TAS2Rs was compared between all CF vs non-CF patients by one way ANOVA with Bonferonni posttest. *TAS2R1* (encoding T2R1) and TAS2R45 (encoding T2R45) were the highest expressed TAS2Rs in this data set and were slightly but significantly decreased in CF patients when compared in this manner. No other T2Rs were significantly different, including T2R14 (shown in two groups as it was included in both arrays. All together, these data support conclusions in the main text that there are minor, if any, differences in TAS2R expression between CF and non-CF nasal epithelial cells.

### Supplementary Figure 2

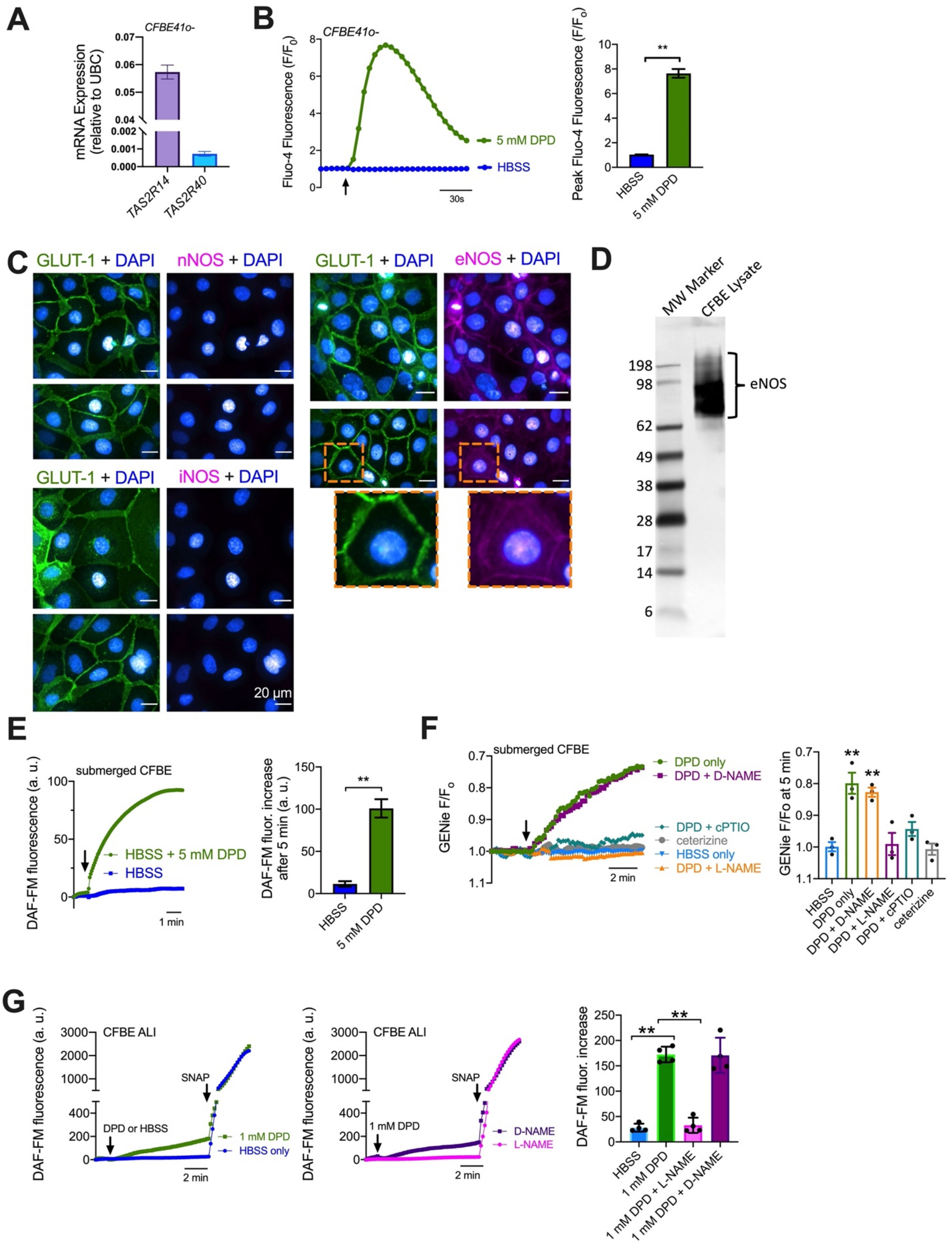

**Supplementary Figure 2: Diphenhydramine (DPD) responses in CFBE41o- (CFBE) cells. (A)** qPCR revealed higher expression of *TAS2R14* (encoding T2R14) compared with *TAS2R40* (encoding T2R40) in CFBE cells, fitting with observations of primary nasal (this study) and bronchial cells [2]. **(B)** 5 mM DPD activated $Ca^{2+}$ elevations (measured by Fluo-4) in submerged CFBE cells. Hank's balanced salt solution (HBSS) only was added as control. Bar graph shows mean ± SEM of n = 3 independent experiments analyzed by Student's *t* test; **p<0.01. (C) Immunofluorescence images of GLUT-1 (membrane marker) plus co-staining for nNOS, iNOS, or eNOS in submerged CFBEs. Representative images shown are taken at the same microscope settings, suggesting membrane staining for eNOS but not nNOS or iNOS. Note that eNOS is palmitoylated and often localized either to the plasma membrane and/or Golgi [3, 4]. We previously observed Golgi localization of eNOS in A549 cells with this antibody [5]. Plasma membrane localization in CFBE cells may have implications for eNOS activation that are beyond the scope of this current study. These data do, however, support expression of eNOS in CFBEs. **(D)** Western blot analysis revealed eNOS expression in CFBE cells. Western blotting was carreid out as described [2]. Briefly, cultured CFBE cells were scrapped in PBS then re-suspended in RIPA buffer followed by lysis via sonicating water bath for 4 rounds of 30s pulses. After centrifugation at 800 x *g* for 8 min at 4°C, the supernatant was collected, and protein concentration estimated via BioRad DC Protein Assay (Hercules, CA). Molecular weight marker SeeBlue Plus2 (Thermo Fisher) and 60 μg of cell extract was separated across a 4-12% NuPAGE Bis-Tris gel (Invitrogen) then transferred to a nitrocellulose membrane for subsequent western blot analysis using an eNOS/NOS Type III primary antibody (Fisher Scientific, Cat #610296) and an anti-mouse secondary antibody (Cell Signaling Cat #7076) relative to. Together with IF, Western data also supports expression of eNOS in CFBE cells. **(E)** Submerged CFBE cells exhibited increases in DAF-FM fluorescence (likely reflecting NO production) in response to 5 mM DPD. Traces are representative experiments and bar graph shows mean ± SEM of 4 independent experiments analyzed by Student's t test; ***p*<0.01, HBSS only added as control. **(F)** Submerged CFBE cells were transduced with BacMam encoding Green GENie cGMP biosensor (Montana Molecular, Bozeman, MT), previously used to measure cGMP in airway cells [2] and macrophages [6]. This biosensor gets dimmer as cGMP increases. We observed a decrease in $F/F_o$ reflective of an increase in cGMP (plotted inversely so it the trace tracks cGMP). The increase in cGMP was inhibited by NO scavenger cPTIO (Cayman Chemical) or NOS inhibitor L-NAME (10 μM; 30 min pre-treatment) but not inactive d-NAME. Bar graphs shows change in F/Fo after 5 min stimulation. Significance determined by one-way ANOVA with Dunnett's posttest comparing all values to control (HBSS alone). **(G)** CFBE cells grown at ALI exhibited increases in DAF-FM fluorescence in response to 1 mM DPD that were inhibited by NOS inhibitor L-NAME (10 μM; 30 min pre-treatment) but not inactive analogue D-NAME (10 μM; 30 min pre-treatment). Traces are representative experiments and bar graph shows mean ± SEM of 4 independent experiments analyzed by one way ANOVA with Bonferroni posttest; ***p*<0.01. S-Nitroso-N-Acetyl-D,L-Penicillamine (SNAP; 20 μM; Cayman Chemical) was added at the end of each experiment as a non-specific S-nitrosothiol NO donor to validate that the loaded DAF-FM can detect NO production, further supporting that the DAF-FM changes reflect NO production.
